# Interactive data exploration websites for large-scale electrophysiology

**DOI:** 10.1101/2024.06.07.597950

**Authors:** Daniel Birman, Gaelle Chapuis, Mayo Faulkner, Cyrille Rossant, the International Brain Laboratory, Julius Benson, Joana A Catarino, Anne K Churchland, Fei Hu, Julia M Huntenburg, Anup Khanal, Christopher Krasniak, Petrina Y P Lau, Guido T Meijer, Nathaniel J Miska, Jean-Paul Noel, Alejandro Pan-Vazquez, Noam Roth, Michael Schartner, Karolina Z Socha, Nicholas A Steinmetz, Anne E Urai, Miles J Wells, Steven J West, Olivier Winter

## Abstract

Methodological advances in neuroscience have enabled the collection of massive datasets which demand innovative approaches for scientific communication. Existing platforms for data storage lack intuitive tools for data exploration, limiting our ability to interact effectively with these brain-wide datasets. We introduce two public websites: (Data and Atlas) developed for the International Brain Laboratory which provide access to millions of behavioral trials and hundreds of thousands of individual neurons. These interfaces allow users to discover both the raw and processed brain-wide data released by the IBL at the scale of the whole brain, individual sessions, trials, and neurons. By hosting these data interfaces as websites they are available cross-platform with no installation. By releasing each site’s code as a modular open-source framework, other researchers can easily develop their own web interfaces and explore their own data. As neuroscience datasets continue to expand, customizable web interfaces offer a glimpse into a future of streamlined data exploration and act as blueprints for future tools.

## Introduction

Modern systems neuroscience requires a new approach to curating, exploring, and sharing datasets and results. Existing websites for storing systems neuroscience data ([DANDI](https://www.dandiarchive.org/), [CRCNS](https://crcns.org/), [OpenNeuro](https://openneuro.org/), [Figshare](https://figshare.com/), [Dryad](https://datadryad.org/stash), [Zenodo](https://zenodo.org/), Foster and Deardorff (2017) and Poldrack et al. (2013)) are focused on data storage. Some websites provide additional interfaces for exploring metadata (e.g. Neurosift), Magland et al. (2024) but stop short of allowing access to raw data files. For all of these data archives, exploring an experimental dataset through an intuitive interface would provide a substantial improvement in the experience of researchers seeking to explore and understand experimental results. Developing such intuitive interfaces for systems neuroscience is also an opportunity to encourage data standardization and experimental rigor. As data acquisition efforts continue to grow to brain-wide scales, our ability to interact with and leverage large-scale data sets for analysis depends on the existence of tools for exploration and sharing.

Here we present two websites developed by the International Brain Laboratory for exploring raw data and visualizing the final results of brain-wide analyses. These websites are publicly accessible, intuitive to explore, and act as portals into millions of trials of behavioral data (International Brain Laboratory et al., 2021) and hundreds of thousands of individual neurons (International Brain Laboratory et al., 2023). The websites are a companion to standard data access tools, such as the IBL’s Open Neurophysiology Environment [application programming interface](https://int-brain-lab.github.io/iblenv/) which provide direct access to raw data. By developing these websites as open-source modular frameworks, external users can build their own versions and upload their own data. Our goal is for these websites to be as useful for expert researchers as they are for students learning about neuroscience for the first time.

## Results

We report here the design and architecture of two websites used to interface with the data sets collected by the International Brain Laboratory (IBL) (Abbott et al., 2017). The [*Data* website](https://viz.internationalbrainlab.org/) is used by researchers to explore data sets after acquisition and basic preprocessing (Fig. 1). This website provides a standardized interface to search for a session and then explore data within each session at the level of the full session, individual trials, or neuron clusters. The [*Atlas* website](https://atlas.internationalbrainlab.org/?alias=bwm_feedback) is used by researchers to explore the results of brain-wide analyses, aggregated across regions (Fig. 2). This second website provides an interactive interface with 2D, 3D, and flattened views of the mouse brain for displaying results that span multiple brain regions. Both websites were developed using a modular architecture which makes it possible for external users to develop custom versions of either site.

**Figure 1:**
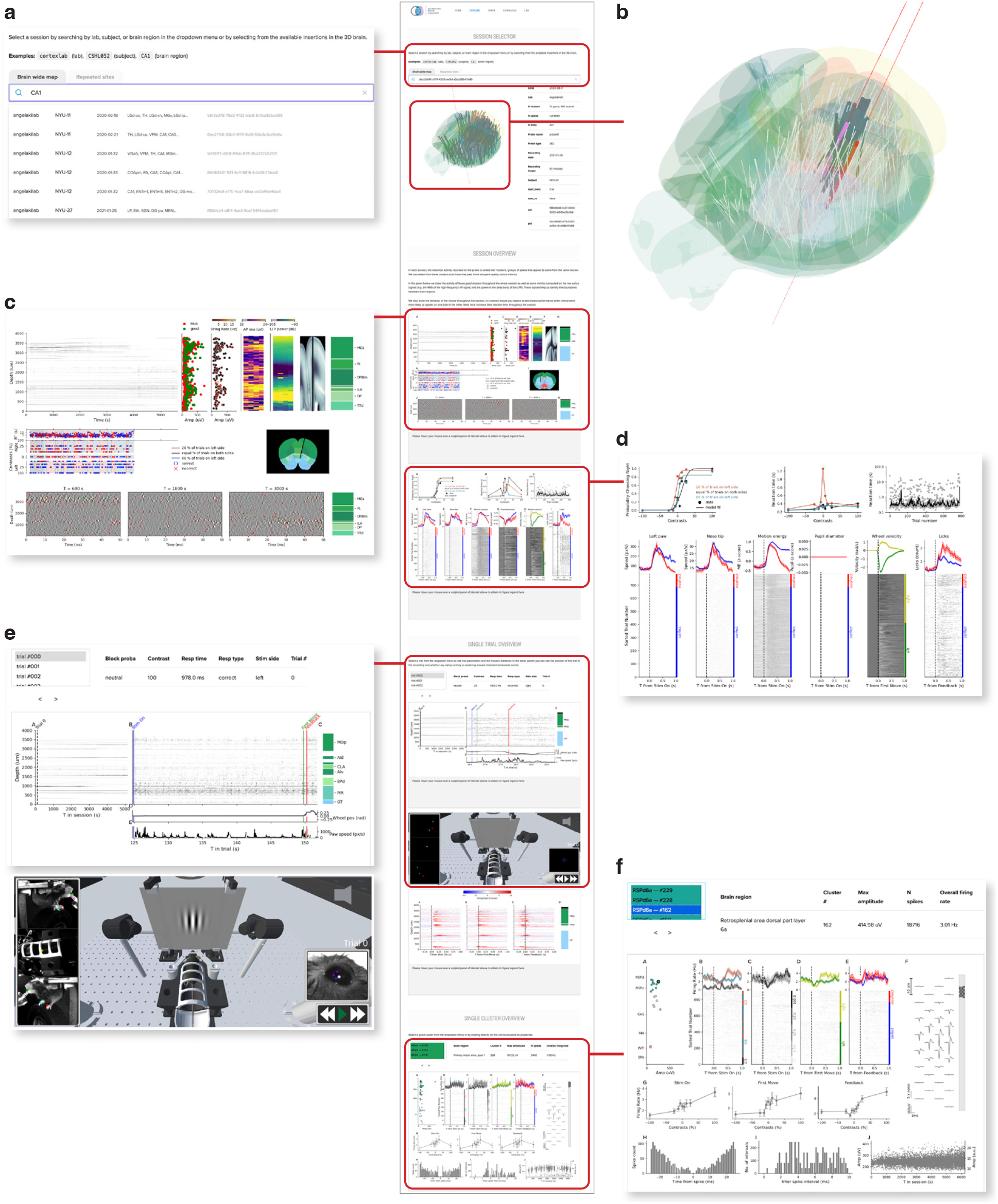
Data visualization website (link). (a) The top section of the Data website, showing the search bar. Users can search by insertion ID, brain region, lab, or subject ID. (b) The 3D brain allows users to hover over insertions with their mouse and click to select them. (c) An example of a session-level figure, showing an overview of the raw electrophysiology data before and after spike sorting, as well as some basic quality control metrics. (d) A second example of a session-level figure, summarizing the behavioral performance of the mouse on the task. (e) An example of a trial-level figure, showing the data from the first trial, including a snippet of the spike raster and wheel movements. The lower section shows a 3D reconstruction of the behavioral rig with annotated videos, on the website this animation can be used to replay the session. (f) An example of a cluster-level figure, here showing rasters and histograms relative to different trial events as well as the spike waveforms on the probe and various quality-control metrics. Not visible in the figure is the “Share” button, that creates a URL link that returns to the same configuration of the website, for easy sharing of a particular example of raw data among researchers.

**Figure 2:**
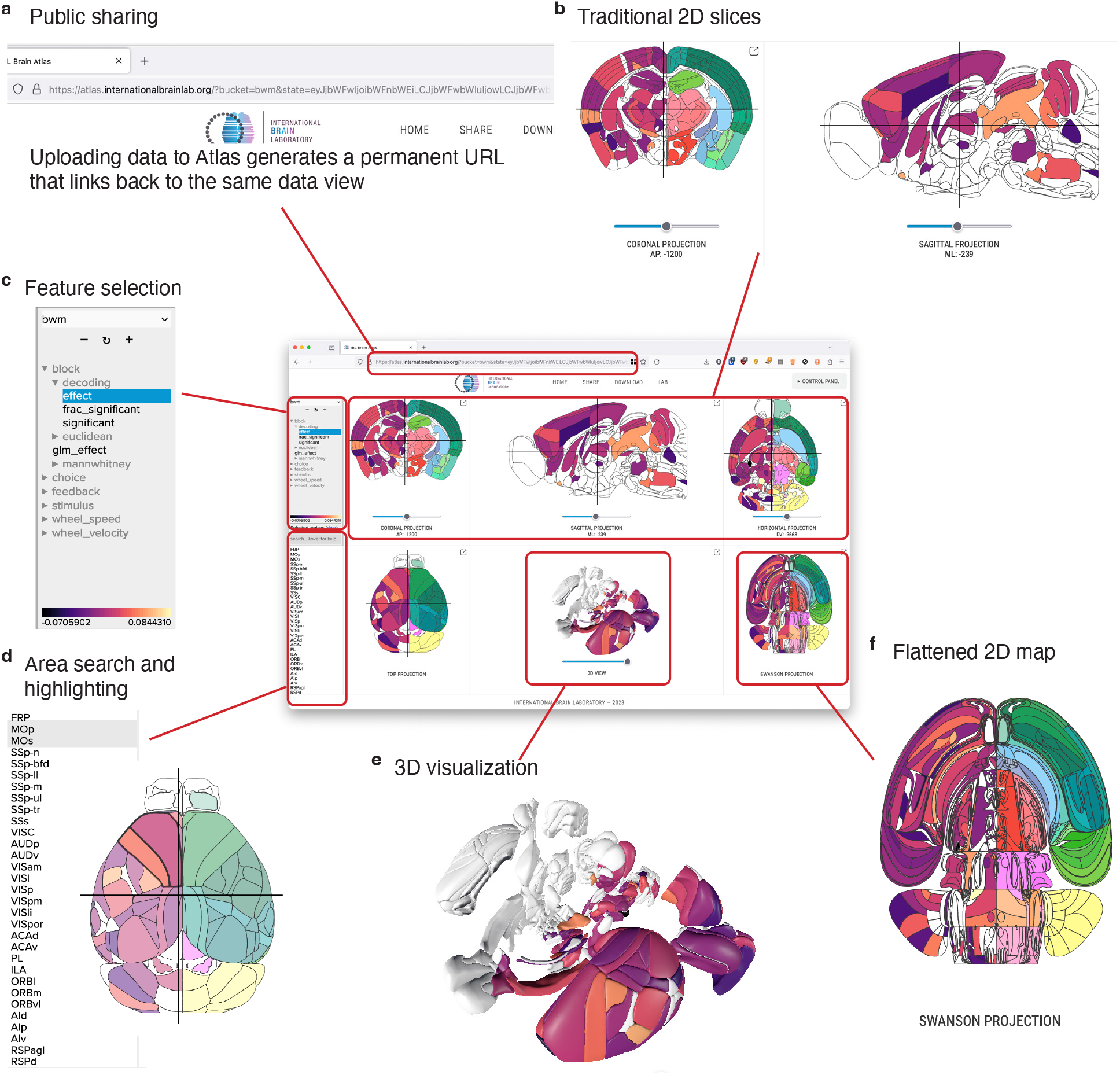
Atlas visualization website (link). (a) The Atlas website generates unique links for each uploaded dataset, allowing users to share their uploaded results. (b) The three upper panels on the site show coronal, sagittal, and axial views of the brain. (c) A dataset selection window lets users choose which “feature” they want to display from the current dataset. (d) Individual areas can be highlighted from a search bar, or by clicking each region in any of the visualization panels. (e) A 3D reconstruction of the brain is shown in the center of the bottom panels. (f) The Swanson flatmap is shown, providing an overview of all regions in the mouse brain at once.

**Figure 3:**
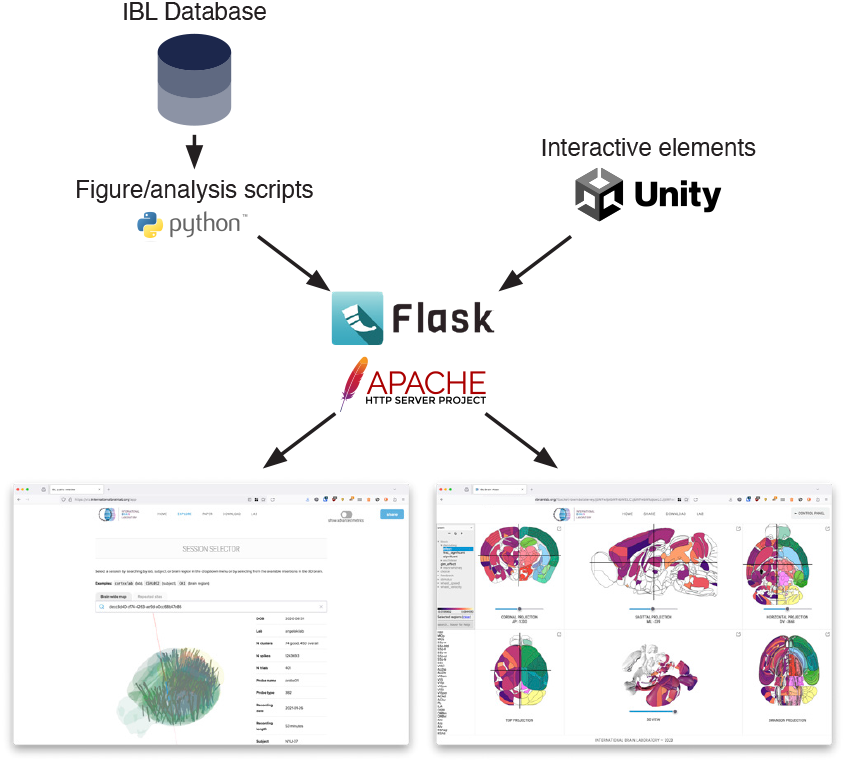
Server-side architecture. The Data and Atlas websites are served by Apache HTTP servers, using Flask to expose a REST API for access to datasets and static images. The figures and data are produced from custom Python scripts, developed with researchers from the International Brain Laboratory. The interactive 3D elements are developed with the Unity Real-Time Development Platform.

### Data website

The IBL data website (https://viz.internationalbrainlab.org/app, Fig. 1) acts as a portal into electrophysiology data sets recorded at brain-wide scale. The site features four main features: search tools and metadata (Fig. 1a,b), per-session figures (Fig. 1c,d), per-trial figures (Fig. 1e), and per-neuron figures (Fig. 1f). The search bar and metadata table allow users to quickly identify an electrophysiology insertion that they want to explore from all the insertions that were performed during an experiment. Sessions can be optionally separated into groups using tabs above the search bar, for example to organize data by publication.

The session, trial, and neuron figures (Fig. 1b-e) each include a variety of panels providing quality control metrics, information about mouse behavior, or electrophysiology. These figures cover the wide range of information that researchers need to know about to evaluate whether a session needs to be excluded from further analysis or to identify issues with the hardware or software data acquisition pipelines prior to further analysis. Some of these figures provide simple interactions to help users quickly move through the dataset. For example, the trial figure allows users to click within the session timeline (Fig. 1e, left side) to move from trial to trial, and the neuron figure allows users to select individual clusters by clicking on them (Fig. 1f, top left panel). For the IBL websites, we also developed an interactive 3D model of the IBL rig which can be used to replay animated individual trials (Bottom of panel, Fig. 1e). This visualization helps users who are unfamiliar with the experimental setup. All of the figure panels include captions, which are revealed when users hover over each panel.

For many experiments conducted in the past century, quickly assessing the quality of behavior and raw data has been difficult. In recent years, researchers in electrophysiology have begun to rely on automated quality control metrics (International Brain Laboratory et al., 2022). We developed the Data website to provide researchers with a tool for fast exploration and quality control of data at the level of a full session, individual trials, or individual neurons. With this goal in mind, one of the most powerful features of the Data website is the **share** button, which creates a unique URL that links back to the same session, trial, and cluster. Researchers have used these links to share interesting examples of unique neurons, quality control concerns, and for orienting outside users to specific aspects of the IBL dataset. The power of the data website comes from its ability to let users explore the dataset in a relatively unconstrained manner.

### Atlas website

One of the most challenging aspects of neuroscience data analysis is the need to visualize findings in their anatomical context. We developed the Atlas website to solve this problem and provide an intuitive interface for 2D and 3D data visualization of brain-wide results.

The IBL Atlas website (https://atlas.internationalbrainlab.org, Fig. 2) acts as an interface into processed results from datasets recorded from mice at brain-wide scale. The website can visualize results at three scales: perneuron, pervoxel, and per-area. Datasets at these different scales are then mapped onto 2D slices (Fig. 2b), flatmaps (Fig. 2f), and 3D reconstructions which can be interactively explored (Fig. 2e). A feature hierarchy allows users to quickly switch the active data in the visualization (Fig. 2d) and a search bar lets users select a subset of regions to highlight (Fig. 2d).

The Atlas website can be used as an interactive figure, allowing static publications to link to dynamic versions of their results. We designed permanent links for each of the major results in the IBL Brain-wide platform project(International Brain Laboratory et al., 2023), so that readers could explore the dataset on their own terms alongside the static figures included in the publication. Users can create their own permanent links using a **share** button to return to a particular combination of dataset, selected feature, and selected areas (Fig. 2a).

### Customizing websites

Both the Data and Atlas websites can be customized to display data from experiments outside of the IBL. For the Data website, we provide a [generic template](https://github.com/int-brain-lab/website/tree/generic). To use the template, an experiment needs to have a concept of a “session”, “trial”, and individual “unit” or cluster of neurons. Users format their experiment metadata according to the specification and place their per-session, per-trial, and per-cluster figures into the folder hierarchy. The template then handles swapping figure images according to the selected session, trial, and cluster. A [tutorial page](https://github.com/int-brain-lab/website/blob/generic/README.md) with additional instructions can be found in the repository.

For the Atlas website, we developed an API that allow users to upload their personal data to our server. Users can install the API using ‘pip install iblbrainviewer’ into a Python environment, and instructions and tutorials are available [online](https://github.com/int-brain-lab/iblbrainviewer/tree/main/docs). Data can be uploaded formatted as points, volumes, or per-area tables. Users can also launch their own [Atlas server](https://github.com/int-brain-lab/ephys-atlas-web) to deploy the website on their own hardware.

## Discussion

We described here two visualization websites that act as interfaces to large-scale electrophysiology data at massive brain-wide scale. These websites provide deep access to the IBL Brain-wide Map (International Brain Laboratory et al., 2023) and are developed as open-source tools for reuse by the community.

Our Data and Atlas websites are part of a larger trend in neuroscience of releasing large-scale datasets with companion visualization tools. These companion tools are often much more capable than the data browsers available in a typical dataset archive which often expose metadata but not in an intuitive or interactive manner. Companion tools for data visualization vary widely in their scope and depth but all share the ability for users to explore and discover aspects of a data release on their own. An early example of this kind of tool was included with the Allen Common Coordinate Framework (CCF, Wang et al. (2020)) which includes an [interactive 3D Viewer](https://connectivity.brain-map.org/3d-viewer) where users can explore the 3D atlas alongside connectivity and tracing data from across the mouse brain. The European EBRAINS initiative takes a similar approach, with 3D explorers available for reference atlases (https://atlases.ebrains.eu/viewer/). Custom websites are also commonly developed to explore specific aspects of a dataset, such as the brain observatory website for the Allen Brain Atlas Visual Coding dataset (https://observatory.brain-map.org/visualcoding) which affords users tools to examine the visual coding properties of individual units. There are also general-purpose rendering tools intended for neuroscientists which have been used by large-scale data releases to support data exploration, for example Neuroglancer (https://github.com/google/neuroglancer), brainrender (Claudi et al., 2021), and Urchin (https://github.com/VirtualBrainLab/Urchin) each of which support volume rendering, 3D meshes, and other neuroscience-specific feature visualization. Neuroglancer has been used by large-scale data releases such as MICrONS (Consortium et al., 2021) to support data exploration. Urchin has been used to create custom web portals for smaller scale projects such as Neuropixels Ultra (Ye et al., 2023) as well as the websites described here.

While exploratory interfaces now exist for a handful of datasets, the expertise and funding necessary to develop these tools remains expensive and most datasets, including many large-scale ones, remain accessible only as raw or processed data files hosted on archiving services (de Vries et al., 2023). By developing powerful exploratory tools as open-source generic frameworks, our intent is that the Data and Atlas websites act as both a benchmark for future large-scale dataset releases and blueprints for future data exploration tools.

## Methods

We report here the design and architecture of two websites developed to allow exploration of large-scale neuroscience data. Details about the acquisition, quality control, preprocessing, and analysis of the data sets can be found in their respective papers: for the behavioral task (International Brain Laboratory et al., 2021), reproducible electrophysiology dataset (International Brain Laboratory et al., 2022) and brain-wide map dataset (International Brain Laboratory et al., 2023).

### Data visualization website

The Data website is an HTML/CSS/Javascript page which serves static PNG images from a Flask application.

The Data website has four main sections: a search bar and 3D selector which allow users to choose an insertion of interest, a per-session figure, per-trial figure, and per-cluster figure. The search bar performs a simple lookup, matching for area acronyms (e.g. VISp), unique session identifiers, and mouse metadata (e.g. mouse names) using the [algolia](https://github.com/algolia/autocomplete) library. To generate the figures, custom scripts in Python were created using the [Open Neurophysiology Environment](https://int-brain-lab.github.io/ONE/). For each session, trial, and cluster in the original datasets (International Brain Laboratory et al., 2022; International Brain Laboratory et al., 2023) figures were generated using the raw or pre-processed data. These figures were placed in a folder hierarchy for easy access from the server. At run-time, the website swaps the PNG images on the client-side according to the requested session, trial, and cluster, respectively. The 3D selector and trial viewer elements were created using Urchin (https://github.com/VirtualBrainLab/Urchin) using the Unity Real-Time Development Platform and exported for WebGL.

To allow for mouse-click interaction with the images, we stored the pixel coordinates of each trial or cluster. For trials, we stored in the x-axis position of the trials in time for the full session timeline or the x-axis position of the start and end of each individual trial for the individual trial timeline. By saving these data, the page can accept mouse clicks, process the position of the click, and then update the current trial according to which trial is closest in time to the position of the click. The same general logic was used to allow users to select clusters by clicking on the cluster figure.

The two interactive 3D elements built in Unity (the 3D brain showing the insertions and the “trial viewer” reconstruction) also communicate bidirectionally with the Javascript elements. Each system sends a serialized message to update the current selected insertion or trial, respectively, when these events are triggered. For the trial viewer, we also had to handle continuous time to synchronize the Javascript figure (which shows the current time as a vertical red bar) with the 3D reconstruction which replays the videos. To simplify video synchronization, we concatenated the four videos into a single video, which is cropped into the different displays visible in the scene. The frame time of the video is translated into the current real time in the experiment, and this information is then sent to the Javascript code to update the figure. Clicking on the figure to select the current time sends a serialized message to the Unity elements which moves the videos to the corresponding frame.

### Atlas website

The Atlas website is an HTML/javascript page developed with custom Javascript code and hosted by an Apache server (https://httpd.apache.org/). To handle large data files we use Pako and gzip for compression. We use Normalize.css to create the layout and styling.

The back-end for the Atlas website is a public Flask server which exposes a REST API. The REST API (1) serves static datasets which include features at the level of areas, clusters, and volumes, and (2) handles receiving uploads of that data and user authentication. Data is stored on the server in static JSON files with a basic folder structure. At upload, users define a folder name and the server copies this information in metadata files. Additional features are then stored as subfolders. To authenticate users, each folder has a private token stored securely on the server and on the computer used to create the folder, these tokens can be shared to allow uploads from multiple locations. A master token is stored on the server and must be provided by users to create new folders.

Visualizations were generated in two pipelines. The 2D area visualizations were generated by converting the CCF atlas (Wang et al., 2020) annotations into path boundaries in the SVG vector image format. These were simplified using Ramer-Douglas-Peucker Algorithm, further simplified through Inkscape (via the command-line interface, parameter 0.0007), and finally cleaned with the svgo package and a custom Python script. 2D volume and slice visualizations were created by simply looping over points and applying these to the pixels of an HTML “canvas” element. The Swanson flatmap (Hahn et al., 2021) was converted to the SVG image format for use on the website. The 3D visualizations were created by integrating the Urchin application (https://github.com/VirtualBrainLab/Urchin/) and using Urchin’s Javascript API to control the visibility of areas, particles, and volumes in the 3D scene.

## Acknowledgments

We extend our thanks to all International Brain Laboratory members and alumni for their feedback and support on these projects, in particular Liam Paninski for significant feedback on the design of the websites and Tatiana Engel for manuscript comments. We acknowledge the support of the Washington Research Foundation Postdoctoral Fellowship to DB, as well as the Wellcome Trust (216324), Simons Foundation, and NIH (U19NS123716) to the IBL.

## Contributions

DB - conceptualization, resources, software, visualization, writing - original draft & editing. GC - conceptualization, resources, data curation, supervision, writing - editing. MF - conceptualization, resources, data curation, software, visualization, writing - editing. CR - conceptualization, resources, software, visualization, writing - editing. JB, JC, FH, AK, CK, PL, GM, NM, JPN, APV, NR, KS, AE - investigation, JH, MS, MW, SW, OW - resources, data curation, software, AC, NS - conceptualization, supervision, funding. Authors are listed in alphabetical order, not by contribution.

